# NaV1.1 and NaV1.6 selective compounds reduce the behavior phenotype in a novel zebrafish model for Dravet Syndrome

**DOI:** 10.1101/675082

**Authors:** Wout J. Weuring, Sakshi Singh, Linda Volkers, Martin Rook, Ruben H. van ‘t Slot, Marjolein Bosma, Marco Inserra, Irina Vetter, Nanda M. Verhoeven-Duif, Kees P.J. Braun, Mirko Rivara, Bobby P. C. Koeleman

**Author notes:** **Correspondence to** B. P. C. Koeleman, Department of Genetics, Centre of Molecular Medicine (CMM), University Medical Centre (UMC), STR.1.305, Lundlaan 6, 3584 CG, Utrecht, The Netherlands.

## Abstract

Dravet syndrome is caused by dominant loss-of-function mutations in SCN1A which cause reduced activity of Nav1.1 leading to lack of neuronal inhibition. On the other hand, gain-of-function mutations in SCN8A can lead to a severe epileptic encephalopathy subtype by over activating Na_V_1.6 channels. These observations suggest that Nav1.1 and Nav1.6 represent two opposing sides of the neuronal balance between inhibition and activation. Here, we hypothesize that Dravet syndrome may be treated by either enhancing Nav1.1 or reducing Nav1.6 activity. To test this hypothesis we generated and characterized a novel DS zebrafish model and tested new compounds that selectively activate or inhibit the human Na_V_1.1 or Na_V_1.6 channel respectively. We used CRISPR/Cas9 to generate two separate *Scn1Lab* knockout lines as an alternative to previous knock-down models. Using an optimized locomotor assay, spontaneous burst movements were detected that were unique to Scn1Lab knockouts and disappear when introducing human SCN1A mRNA. Besides the behavioral phenotype, Scn1Lab knockouts show sudden, electrical discharges in the brain that indicate epileptic seizures in zebrafish. Scn1Lab knockouts showed increased sensitivity to the convulsant pentylenetetrazole and a reduction in whole organism GABA levels. Drug screenings further validated a Dravet syndrome phenotype. We tested the Na_V_1.1 activator AA43279 and our newly synthesized Na_V_1.6 inhibitors MV1369 and MV1312 in the Scn1Lab knockouts. Both type of compounds significantly reduced the number of burst movements. Our results show that selective inhibition of Na_V_1.6 could be just as efficient as selective activation of Na_V_1.1 and these approaches could prove to be novel potential treatment strategies for Dravet syndrome and other (genetic) epilepsies. Compounds tested in zebrafish however, should always be further validated in other model systems, preferably human derived.

## Introduction

Dravet syndrome (DS), previously known as severe myoclonic epilepsy of infancy (SMEI), is a severe form of epilepsy for which current medication strategies remain largely inefficient. Promising new drugs that act on the serotonin pathway such as Fenfluramine (FA), show efficacy in reducing seizures in 50% to 90% of the patients ^[1]^. However, drug side effects may still limit their use, underscoring the need for further drug discovery.

Of all DS patients, 95% carry a *de novo* heterozygous mutation in *SCN1A* ^[2]^, which encodes the pore forming α-subunit of neuronal voltage gated sodium channel (VGSC) type 1 (Na_V_1.1). Na_V_1.1 ion channels are the primary Na^+^ channels in GABAergic interneurons ^[3] [4]^ and induce the fast depolarization of neuronal membranes during action potential initiation. The majority of *SCN1A* mutations in DS are loss-of-function (LOF) mutations resulting in dysfunctional Na_V_1.1 channels, or reduced Na_V_1.1 expression ^[5] [6]^. Consequently, excitability and action potential amplitude of interneurons are attenuated leading to reduced GABA release ^[40]^.

Another brain VGSC subtype, Na_V_1.6 is one of the two main sodium channels expressed in pyramidal neurons, which are responsible for excitatory signals via glutamate excretion ^[7]^. *SCN8A*, which encodes the Na_V_1.6 α-subunit is also related to epilepsy and approximately 100 mutations have been reported in patients with severe Early Infantile Epileptic Encephalopathy subtype 13 (EIEE13). Unlike the clear LOF mutations in *SCN1A*, the majority of the functionally tested *SCN8A* mutations result in gain-of-function (GOF) of Na_V_1.6 ^[8]^. GOF mutations in Na_V_1.6 cause channel hyperactivity due to augmented excitability and firing rates of pyramidal cells concurrent with an increase in glutamate release.

This disease mechanism is reflected by the therapeutic response of VGSC blockers. Various clinical reports have shown that *SCN8A*-related epilepsy patients benefit from VGSC blockers ^[9] [10]^, contrasting their inefficacy, or even detrimental effects in DS ^[11]^. These observations indicate that Na_V_1.1 and Na_V_1.6 represent two opposing sides of the neuronal balance between inhibition and activation. We hypothesize that LOF mutations in *SCN1A* have a major negative effect on neuronal inhibition via hypo-activity of inhibitory interneurons, shifting the balance to neuronal hyperactivity. In contrast, GOF *SCN8A* mutations cause increased activity of excitatory pyramidal neurons, also shifting the VGSC-related balance towards neuronal hyperactivity. This model suggests that either selective activation of Na_V_1.1 or selective inhibition of Na_V_1.6 could be a therapeutic approach in the treatment of both DS and *SCN8A*-related epilepsy.

The therapeutic effect of Na_V_1.1 activation was previously shown in DS mice using spider venom peptide Hm1a (HmTx1) that led to a reduction of seizures and mortality ^[12]^. In another study using a different mouse strain, Hm1a was found to be lethal at picomolar doses within two hours ^[13]^, indicating that safety and administration needs to be further studied. Another Na_V_1.1 activator is the chemically synthesized small molecule AA43279, which showed anti-convulsant properties *in-vivo* but has not been tested in an animal model for DS. In comparison with Hm1a, AA43279 is less selective for Na_V_1.1, indicating it could activate other Na_V_ subtypes at lower concentrations. Nevertheless, AA43279 did not show lethality or sedative and ataxic side-effects at a concentration of 300mg/kg in mice ^[15]^. Inhibition of Na_V_1.6 in DS was previously mimicked by introducing an *SCN8A* missense mutation in DS mice, which reduced seizure susceptibility and increased their lifespan ^[14]^. Compounds that block Na_V_1.6 selectively have not been published to date. To test if both Na_V_1.1 agonists and Na_V_1.6 antagonists could be beneficial in the treatment of DS we generated a novel DS zebrafish model by knocking out the *Scn1Lab* gene using CRISPR/Cas9. In humans, most of the mutations that cause DS are severe truncating mutations, while mild missense mutations are observed in patients with a milder epileptic phenotype ^[41]^. To mimic the genetic architecture of DS in human patients as best possible, the animal model should display a 50% haploinsufficiency of *SCN1A.* Zebrafish likely carry two orthologues for the human *SCN1A* gene; *Scn1Laa* and *Scn1Lab*. While the expression of these genes does not overlap at embryonic-but only at larval stages ^[18] [25]^, they have a shared functional role in epilepsy ^[20]^. Since 2010, three other *Scn1Lab* zebrafish models have been introduced, generated by N-ethyl-N-nitrosourea (ENU) mutagenesis ^[18] [19]^ or morpholino antisense oligomers (MO) ^[17]^. All three display an epileptic phenotype that includes hyperactivity and epileptic spikes recorded from the brain. Nevertheless, CRISPR/Cas9 is currently the most efficient technique to specifically disrupt the gene that is targeted, unlike ENU mutagenesis ^[21]^, and acts on the DNA rather than protein as with MO based approaches ^[23][24]^. To mimic DS in human patients, we generated two zebrafish models with different truncating mutations in *Scn1Lab* using CRISPR/Cas9.

The epileptic phenotype and drug response in zebrafish larvae can be measured by quantifying high velocity burst movements which are indicative of epileptic seizures in fish. The effect of anti-epileptic drugs on this unique behavior phenotype was found to be well correlated with actual epileptic spikes recorded from the zebrafish brain ^[33]^. For this reason, we used the local field potential set-up in combination with a behavior essay to establish the initial phenotype in *Scn1Lab* knockouts, but used the locomotor assay as a single read-out on the drugs tested. After further molecular validation of the animal model, we tested Na_V_1.1 agonist AA43279 and synthesized and two novel Na_V_1.6 channel antagonists, MV1369 and MV1312. Our results show that selective targeting of Na_V_1.1 or Na_V_1.6 ion channels reduced seizure activity in *Scn1Lab* knockout zebrafish, indicating a restoration in neuronal signaling.

## Results

### Generation of Scn1Lab knockout zebrafish

Heterozygous and homozygous *Scn1Lab* knockout zebrafish were generated using CRISPR/Cas9. A 13 bp deletion was created in *Scn1Lab* exon 10, generating a STOP codon on position 474, truncating the *Scn1Lab* protein (**S2a-c Figure**). From here on, *Scn1Lab* knockout indicates 5 days post fertilization (dpf) larvae carrying the homozygous 13bp deletion in *Scn1Lab*. A second allele, carrying a 5 bp deletion in exon 10, leading to a STOP codon on position 487 was generated in parallel to confirm the knockout phenotype (**S2b Figure**). To validate Cas9’ specificity in DNA editing, potential off-target regions were sequenced. No off-target editing of Cas9 was observed (**S3 Figure**). To verify the presence of the genomic deletion at transcription level, cDNA was sequenced, resulting in detection of the deletion (**S4 Figure**). *Scn1Lab* knockouts showed a similar morphological phenotype as observed in previous *Scn1Lab* knock-down models ^[17] [18] [19]^ including hyper-pigmentation and the absence of an inflated swim bladder (**Figure 1a**). Heterozygous knockout *Scn1Lab* larvae showed no apparent morphological difference compared to wildtype zebrafish (**Figure 1b-c**). As a control for the morphological differences of *ScnLab* knockouts, we generated *nisb-WT* control zebrafish ^[17]^ that lack an inflated swim bladder (**S5 Figure**).

**Figure 1.**
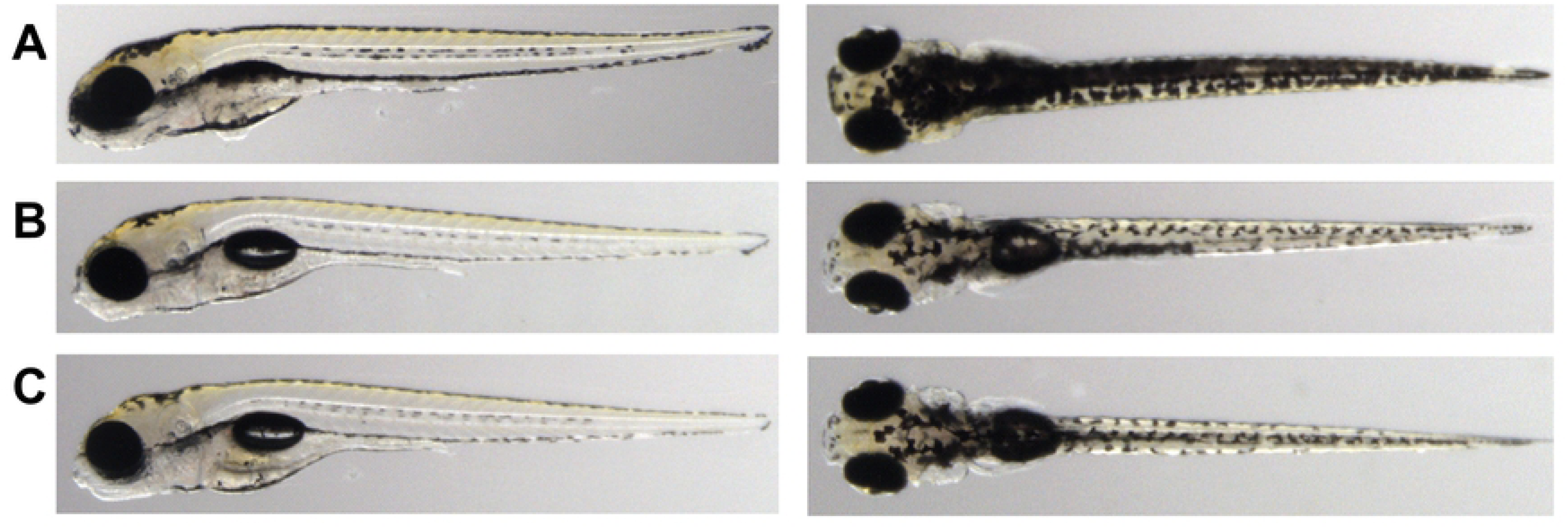
Morphology of Scn1Lab knockout. A) 5 dpf Scn1Lab homozygous knockout larvae show hyperpigmentation and the absence of an inflated swim-bladder. These morphological defects are absent in heterozygous (B) or wildtype (C) zebrafish larvae.

### Epileptic phenotype of Scn1Lab knockouts

*Scn1Lab* knockouts showed behavior comparable to previously described *Scn1Lab* knock-down models ^[17][18][19]^ including hyperactivity and high velocity burst movements (**S6 video**). Using optimized parameters, these specific, high velocity (>50mm/s) burst movements were separated from regular locomotor data to yield a burst movement assay. *Scn1Lab* knockouts showed a significantly higher number of spontaneous burst movements (**Figure 2a**). Using the *nisb-WT* control zebrafish, burst movements were found to be unique to *Scn1Lab* knockouts and not caused by the absence of an inflated swim bladder (**Figure 2b**). No spontaneous burst movements were observed in heterozygous knockout *Scn1Lab* larvae (**Figure 2a**). Using a local field potential (LFP) configuration for zebrafish embryos ^[26]^, abnormal brain activity was observed in Scn1Lab knockouts including spontaneous, single- and poly spiking electrical discharges. The LFP patterns recorded from Scn1Lab knockouts resembles those observed in previous *Scn1Lab* knock-down zebrafish models ^[17] [18]^ (**Figure 2f**).

**Figure 2.**
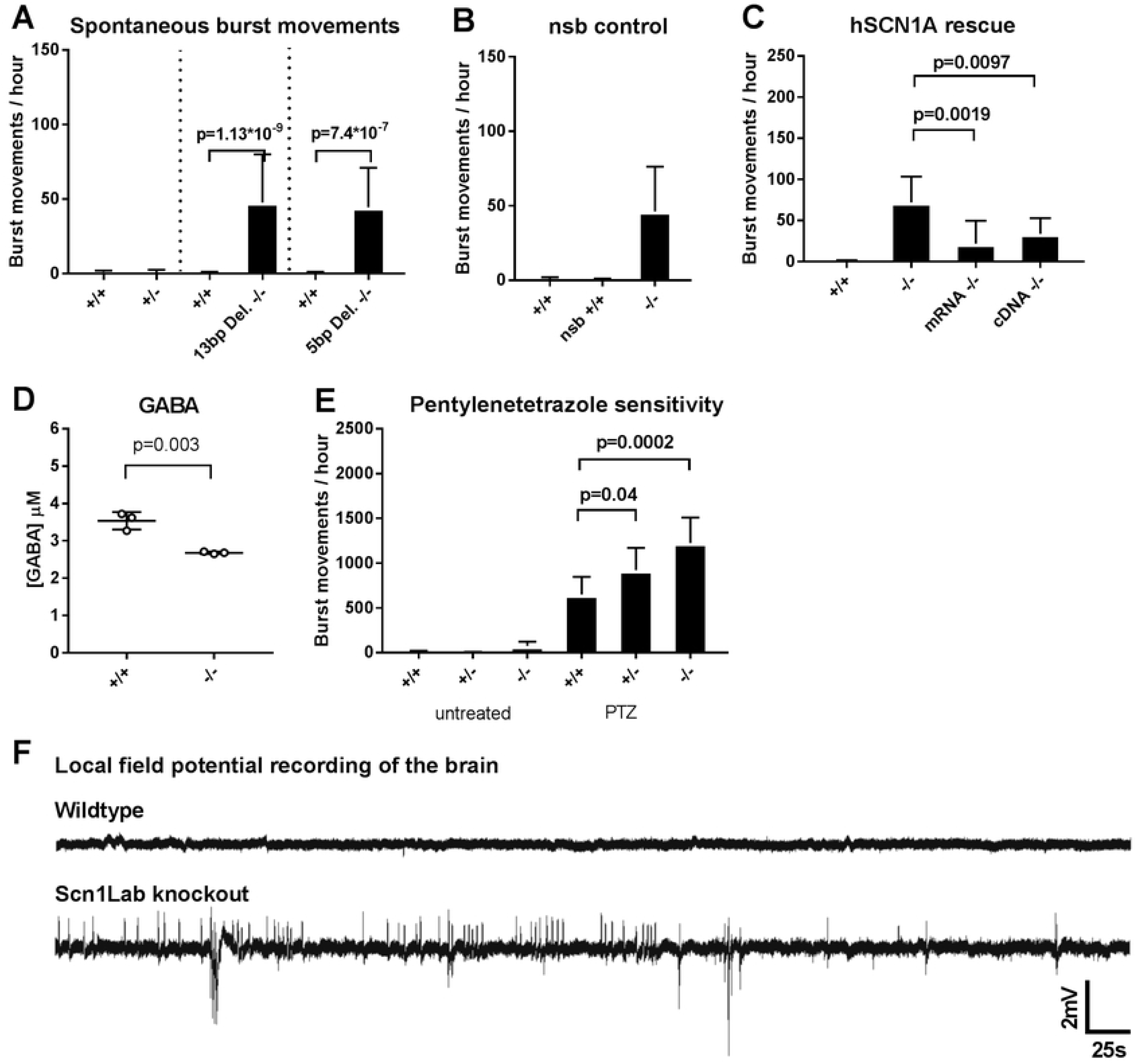
A) Spontaneous burst movements quantified in two knockout lines using the locomotor burst movement assay B) Burst movements are unique to scn1lab knockouts, and not caused by the absence of an inflated swimming bladder C) The burst movement phenotype of scn1lab knockouts is partially rescued when human SCN1A, either mRNA or cDNA is introduced D) Scn1Lab knockouts show a reduction in free GABA levels E) Both heterozygote and homozygous Scn1lab knockouts show sensitivity to exposure of 5mM pentylenetetrazole F) Representative non-invasive local field potential recordings from the brain of wildtype and Scn1Lab knockout zebrafish (n=3). Scn1Lab knockout zebrafish show spontaneous electrical discharges that resemble seizure activity. *Error bars = S.D. (-/-) = Scn1Lab knockout, locomotor assay n=12 per group*

### Partial rescue of Scn1Lab knockout burst movements by human SCN1A

*Scn1Lab* is believed to be one of the two functional orthologues for human *SCN1A*, therefore we tested whether the spontaneous burst movement phenotype of *Scn1Lab* knockouts can be rescued by the introduction of human *SCN1A* in our model. Human *SCN1A* mRNA or cDNA was injected in one-cell stage *Scn1Lab* knockout embryos leading to a partial rescue of burst movements. (**Figure 2c**).

### GABA reduction and sensitivity to pentylenetetrazole (PTZ) in Scn1Lab knockouts

To see if the GABAergic tone is disturbed in our *Scn1Lab* knockouts, levels of free GABA were quantified in whole organism by Mass-spectrometry. *Scn1Lab* knockouts showed a statistically significant reduction of GABA (**Figure 2d**). Thus, as *Scn1Lab* knockouts are GABA deficient, they might be more susceptible to convulsing agents that act on the GABAergic inhibitory pathway. PTZ, a GABA antagonist frequently used as convulsant in animal studies was applied to *Scn1Lab* knockouts. Heterozygous and homozygous *Scn1Lab* knockouts showed a statistically significant increase in burst movements when exposed to 5mM PTZ, compared to wildtype zebrafish (**Figure 2e**).

### Pharmacological validation confirms a DS phenotype

Traditional VGSC blockers are known to be inefficient in *SCN1A* related epilepsies and can even sometimes aggravate seizures in Dravet syndrome, likely due to their lack of Na_V_ subtype specificity. To test if VGSC blockers alter the burst movement phenotype in our *Scn1Lab* knockouts, the anti-epileptic drugs (AEDs) Phenytoin (PTH) and Carbamazepine (CBZ) were applied using a short or long drug exposure time. No reduction in burst movements was observed in the *Scn1Lab* knockouts (**Figure 3a, b**) when exposed to PTH or CBZ, confirming the inefficacy of VGSC blockers in DS. Next we tested the anti-seizure effects of GABA enhancing drugs Valproic acid (sodium valproate, VPA) and Stiripentol (STP), which are effective treatments for recurrent seizures in DS. Seizure-like burst movements were significantly reduced when *Scn1Lab* knockouts were exposed to STP (**Figure 3d**). Interestingly, VPA showed an effect only after long exposure, short exposure did not result in burst movement reduction (**Figure 3c**).

**Figure 3.**
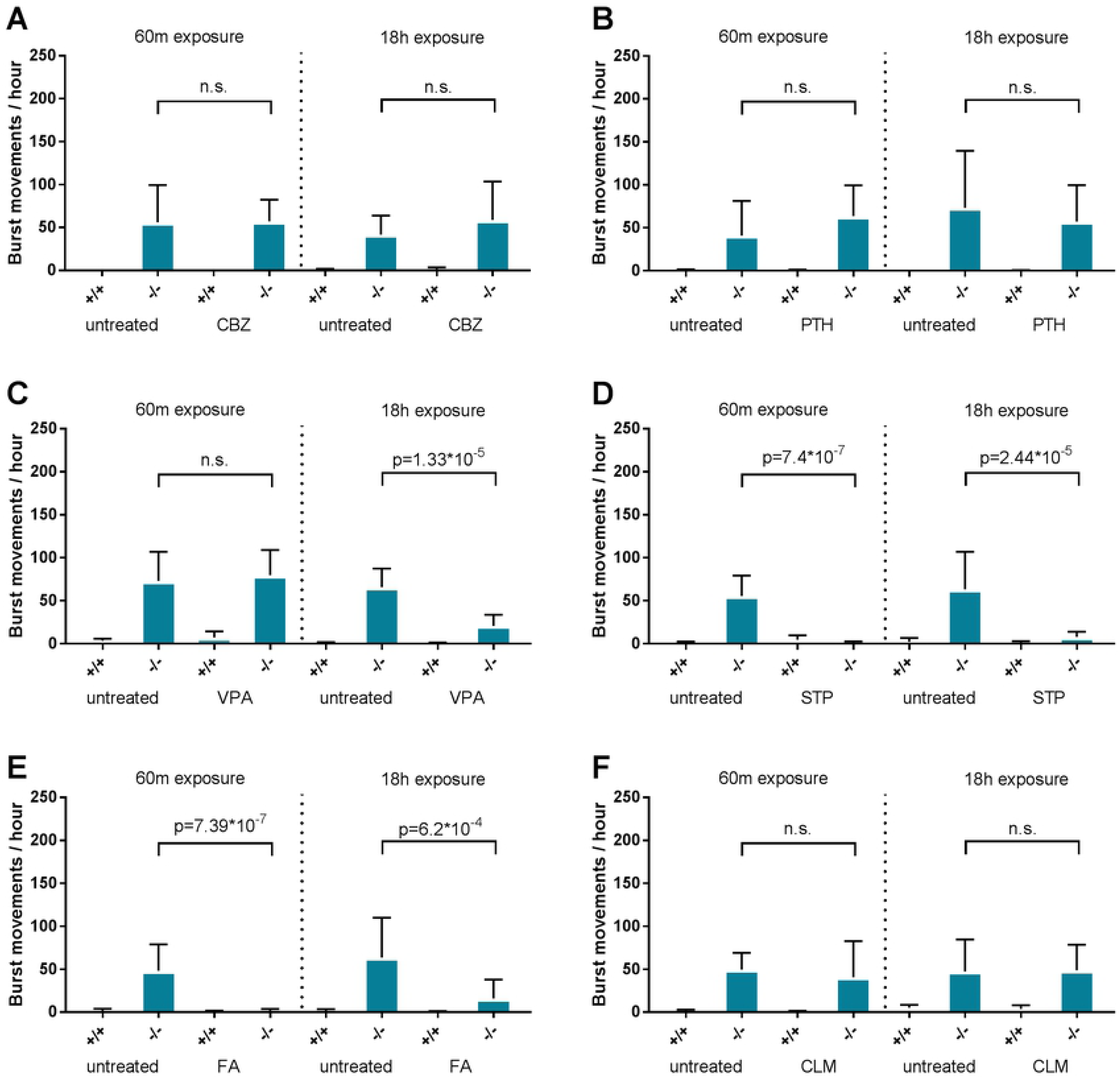
Pharmacologic validation of Scn1Lab knockouts by short (60 minutes) or long (18 hours) exposure of anti-epileptic drugs to the swimming medium A) 50µM Carbamazepine B) 100µM Phenytoin C) 100µM Sodium valproate D) 12.5µM Stiripentol E) 50µM Fenfluramine F) 50µM Clemizole. Dashed lines indicate a novel experimental plate with a seperate experimental group. *Error bars = S.D., (-/-) = Scn1Lab knockout, n=12 per group.*

### Partial seizure reduction by serotonin pathway modulators

Fenfluramine has previously been discovered in the *Scn1Lab* morpholino knock-down model to be effective in reducing DS seizures ^[17]^ and is now shown to be equally effective in our *Scn1Lab* knockout (**Figure 3e**). Clemizole (CLM), an antihistamine that can also bind to the serotonin receptor did not show a significant reduction of burst movements in the *Scn1Lab* knockouts unlike a previous *Scn1Lab* knock-down model ^[18]^. Toxicity of CLM was observed in the *Scn1Lab* knockout long exposure groups, showing mortality and body malformations at previous established concentrations for a short exposure experiment ^[18]^ (**S7 Figure**). When exposed to 50% of this dose, no toxicity was observed and no significant reduction in burst movements was observed (**Figure 3f**).

### Na_V_1.1 selective-but not general VGSC activators reduce seizures

Wildtype zebrafish exposed for a short or long incubation time to the general VGSC activator Veratridine (VRT) developed burst movements, confirming the convulsing effects of VRT in healthy control animals (**Figure 4a**). Interestingly, *Scn1Lab* knockouts revealed no additional increase, nor decrease of burst movements after being exposed to VRT (**Figure 4a)**. When exposed to Na_V_1.1 selective activator AA43279, the number of burst movements in *Scn1Lab* knockouts was significantly decreased (**Figure 4b**). This effect was only observed in the short, but not in the long exposure group, confirming the rapid clearance of AA43279 that has been reported in mice ^[15]^.

**Figure 4.**
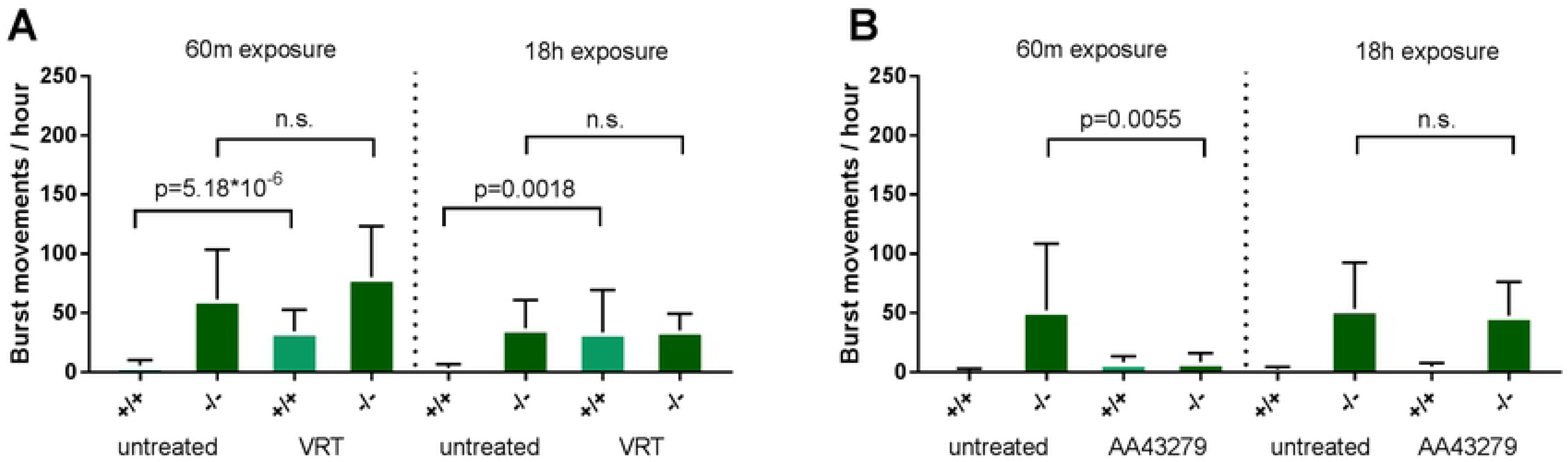
The effect of VGSC activators on burst movements in Scn1Lab knockouts A) 10µM Veratridine B) 5uM AA43279 *Error bars = S.D. (-/-) = Scn1Lab knockout, n=12 per group.*

### Selectivity of MV1312 and MV1369 and efficacy in Scn1Lab knockouts

From a range of unpublished VGSC blocking compounds, several were tested for human Na_V_1.6 selectivity. MV1312 had a 5-6 fold selectivity of Na_V_1.6 over Na_V_1.1-1.7, but comparable blocking affinity for Na_V_1.8 (**Figure 5a**). Na_V_1.8 is a peripheral nervous system ion channel involved in the sensation of pain, and blockage could lead to anti-nociception and pain treatment. As the selectivity over Na_V_1.1 is most important, we chose compound MV1312 to be a suitable candidate for further evaluation in our DS animal model. When exposed to 5µM MV1312, the number of burst movements was statistically significant reduced (**Figure 5b)**, indicating a restoration of neuronal signaling in the epileptic brain. Compound MV1369 showed less selectivity for Na_V_1.6, but did show selectivity over the CNS channel Na_V_1.2 (**Figure 5c**). Interestingly, when applied MV1369, the number of burst movements was also statistically significant reduced (**Figure 5d)**. As the DNA and proteins differ between animal models and humans, it could be that compound selectivity for human proteins is actually not represented in animal models.

**Figure 5.**
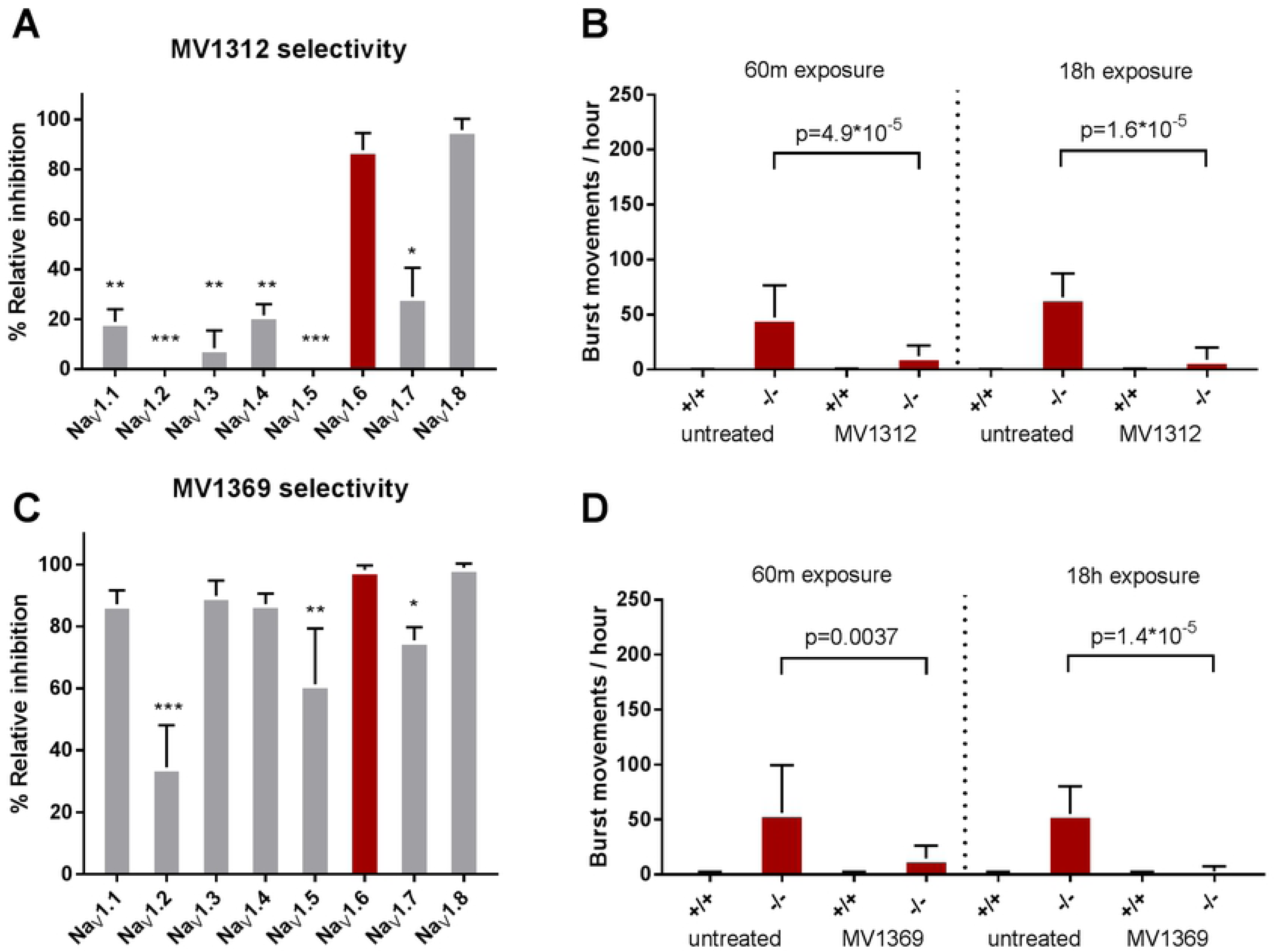
Selectivity of ion channel blocking compounds A) MV1312 show blocking selectivity for Na_V_1.6 over the other ion channel subtypes but not Na_V_1.8. B) The effect of 5µM MV1312 on locomotor burst movements in Scn1Lab knockout zebrafish C) MV1369 shows blocking selectivity for Na_V_1.6 over Na_V_1.2, Na_V_1.5 and Na_V_1.7. D) The effect of 50µM MV1369 on locomotor burst movements in Scn1Lab knockout zebrafish *Error bars = S.D., (-/-) = Scn1Lab knockout, locomotor assays n=12 per group, selectivity assays n=3 per group*, * <*0.05* **<*0.005* ***<*0.0005.*

## Discussion

Here, we present two CRISPR/Cas9 generated knockout zebrafish models for *SCN1A*-related epilepsies, including Dravet syndrome. The phenotype of *Scn1Lab* knockouts is characterized by spontaneous burst movements and sudden electrical discharges in the brain, a phenotype comparable to previous *Scn1Lab* knock-down models. Here, we used an optimized locomotor assay to extract burst movements from regular movement activity and found that the introduction of human *SCN1A* mRNA rescues this phenotype, indicating that *Scn1Lab* and SCN1A have a comparable function. *Scn1Lab* knockouts show a reduction in whole organism GABA levels, and are more sensitive to convulsions induced by GABA antagonist PTZ, compared to wildtype larvae. These results mimic haploinsufficiency of *SCN1A* in humans that likely affect local GABA levels, leading to the seizure susceptibility in DS patients. By applying standard AEDs and DS specific drugs we observed a comparable pharmacological response as in DS patients, but also highlighted differences in drug response in comparison to previous *Scn1Lab* knock-down models. Finally, we show that the Na_V_1.1 activator AA43279 and Na_V_1.6 selective compounds reduced the burst movement phenotype in *Scn1Lab* knockouts, indicating that selective ion channel inhibition or activation could be beneficial in epilepsy.

In our study, the non-selective VGSC activator Veratridine induced burst movements in wildtype zebrafish confirming its convulsing properties and suggesting that it likely has stronger effects on Nav1.6 inducing seizures, rather than increasing inhibition through Nav1.1. However, there was no effect in the *Scn1Lab* knockouts, underlining that treatment of *SCN1A* haploinsufficiency could benefit from Na_V_ activators and the need for VGSC subtype selective compounds. AA43279 is a small molecule with reasonable selectivity for Na_V_1.1 over the other Na_V_ subtypes and showed a seizure reducing effect in our *Scn1Lab* knockouts. However, with lesser affinity AA43279 could also activate other Na_V_ subtypes (off-target activation), potentially leading to unwanted side effects. For example, off-target activation could lead to myotonia (Na_V_1.4), atrial fibrillation and possible cardiac arrest (Na_V_1.5), seizures (Na_V_1.2) or painful neuropathy (Na_V_1.7 and Na_V_1.8). For this reason, only compounds with high selectivity and efficacy at a very low dose are suitable candidates for translation to human patients. On the other hand, with future improved drug delivery systems such as nanoparticle liposomes targeting the CNS specifically, off-target activation in the peripheral nervous system could be limited.

Inhibition, rather than activation of VGSC is a treatment method that might be preferential, especially when inhibition can be targeted to one or few channel subtypes only. Inhibitors of VGSC have been used for decades in epilepsy patients and although current VGSC blocking drugs such as carbamazepine and phenytoin are hardly selective, they have been proven safe in humans. Therefore, to assess whether inhibition of Nav1.6 can be a novel treatment strategy for DS, also applicable for epilepsy caused by SCN8A gain-of function mutations and perhaps also epilepsy in general, we tested two compounds that selectively inhibit Na_V_1.6 channels. MV1312 showed a seizure reducing effect in our *Scn1Lab* knockouts comparable to Fenfluramine and other DS specific drugs. For this reason, we believe that Na_V_1.6 selective inhibitors could be just as efficient as Na_V_1.1 selective activators and are potentially a safer choice in human patients. Another compound, MV1369 was found to be selective for Na_V_1.6 over Na_V_1.2 and reduces the locomotor phenotype in Scn1Lab knockouts as well. There are several reasons why this type of compound is equally effective in the *Scn1Lab* knockout. First, compounds that are selective for human ion channel proteins, could have a different effect when applied to animal models, as the proteins are not identical to those in humans. Only minor differences on nucleotide level could allow, or limit proper binding of compounds in the protein binding site. Second, not all VGSC genes are evolutionary conserved in zebrafish, but only 6 other ion channel genes exist beside *Scn1Laa* and *Scn1Lab*: *Scn4aa/ab, Scn5Laa/ab* and *Scn8aa/ab* ^[25]^. While the function of these genes is not yet fully understood, it is clear that the absence of the full spectrum of Na_V_ subtypes in zebrafish limits measurable side-effects. As *Scn1Lab* is knocked out in our model, the supposed remaining inhibitory neurotransmission is regulated by *Scn1Laa*, highlighted by the efficacy of the Na_V_1.1 activator AA43279. However, it could very well be that *Scn1Laa* is not a complete functional duplicant of *Scn1Lab*, but carries a shared function, and protein structure of *SCN1A, SCN2A, SCN3A* and even *SCN9A* as proposed earlier ^[25]^. Therefore, efficacy, safety and administration routes of compounds tested in zebrafish, should be further studied in a human model system, with human genes and proteins. To further improve treatment of genetic epilepsy, and reach selective activation or inhibition with more potency and selectivity, drugs should be designed that act on the nucleotide level. As disorders such as DS have a genetic cause, their treatment should also act at the genome-or transcriptome level. For this reason we are looking forward to clinically relevant treatments based on small RNAs, CRISPRi and others that act on the *SCN1A* or *SCN8A* gene directly.

## Materials & Methods

### Zebrafish maintenance & ethics statement

Adult zebrafish stocks of TL strain (Tubingen longfin) were maintained at 28.0°C on a 14/10 hour light/dark cycle under standard aquaculture conditions. Embryos were kept under constant light condition in embryo medium (E3); 5 mM NaCl, 0.17 mM KCl, 0.33 mM CaCl2, and 0.33 mM MgSO4) at 28.5°C. For all experiments described, larvae of 5 dpf were used. All zebrafish experiments carried out were approved by the Animal Experimentation Committee of the Royal Netherlands Academy of Arts and Sciences. For imaging, larvae were embedded in 2% low-melting point agarose prepared with E3 medium.

### sgRNA design and Cas9 preparation

Gene specific guide RNAs (sgRNAs) were designed targeting *Scn1Lab* exon 10, using CHOPCHOP ^[29]^ with an off-target binding cut-off of 4 or more base pair mismatches. sgRNA oligonucleotides were synthesized according to previously described methods ^[30]^. Oligos are listed in **S1 Table**. Capped Cas9 mRNA was created by *in vitro* transcription using Thermo Fisher mMESSAGE mMACHINE™ SP6 Transcription Kit from pCS2-nls-zCas9-nls (Addgene#47929)

### CRISPR/Cas9-sgRNA injections and genotyping

Fertilized eggs were injected with 2 nl of a solution containing 500ng *Cas9* mRNA, 150ng sgRNA and 0.2uL Phenol Red. sgRNAs targeting efficiency was tested by PCR amplifying the target region of 8 injected eggs on 2dpf. Primer sequences used for genotyping are listed in **S1 Table**. Injected embryos were raised to adult mosaic fishes. F0 founders were identified from week 10 by genotyping. F0 founders were outcrossed with wildtype fish to generate F1 embryos. F1 embryos were sampled for genotyping to confirm germline transmission of the mutation. The remaining F1 embryos were raised to adulthood and genotyped at week 10 by fin-clipping. Heterozygous knockouts carrying the same mutations were selected and crossed to raise the homozygous knockout F2 generation.

### CRISPR off-target assay

The gene-specific region including the protospacer adjacent motif (PAM) of the sgRNA was submitted to CCtop ^[31]^ to detect potential off-target sites. Five potential off-target sites with a maximum of four mismatches were selected for Sanger sequencing. Target sites, locations and the primers used for sequencing are included in **S1 Table.**

### GABA measurements

*Scn1Lab* knockout or wildtype larvae were pooled (n=20) in eppendorf tubes in triplicates. Samples were centrifuged at 3500 rpm for 12 minutes at 4C° after which they were lysed in 500µL pre-chilled methanol using 0,5 mm zirconium oxide beads in a bullet blender. Samples were diluted 10 times and frozen at −80C° until day of analysis. A detailed Mass-spectrometry procedure be found in the Supplementary data.

### Locomotor assay

Locomotor experiments were performed under dark conditions on 5dpf embryos placed in a flat bottom 48-well cell culture dish filled with 1mL E3/drug solution. Larvae were placed in single wells at 4 dpf to prevent stress from pipetting on the day of measurement. Movements were tracked in an automated tracking device (ZebraBox™ apparatus; Viewpoint, Lyon, France) for 90 minutes, stacked in 10 minute bins, of which the first 30 minutes were removed as habituation time for the locomotor chamber. The final recording time for all locomotor experiments yielded one hour total. Threshold parameters of viewpoint software were freezing 1, sensitivity 8, burst 50 resulting in a burst movement cut-off value of >50mm/s. Locomotor activity was quantified and analyzed by ZebraLab TM software (Viewpoint).

### hSCN1A Rescue

The SCN1A plasmid, which encodes the human neonatal Nav1.1 ion channel, was previously described ^[28][32]^. Capped hSCN1A mRNA was *in vitro* transcribed using Thermo Fisher mMESSAGE mMACHINE™ T3 Transcription Kit. For mRNA injections, 200ng/ul mRNA was injected in the yolk of one-cell stage zebrafish embryos.

### Local field potential recordings

The constructed LFP setup and concurrent recording settings were based on previous work ^[26]^ with slight modifications. In short, a glass electrode, connected to a high-impedance amplifier, was filled with 1mM NaCl. A larva was then embedded in 2% low-melting-point agarose (Invitrogen) and the glass electrode (2–7 MΩ) placed on top of the forebrain of the larva. The recordings were performed in current clamp mode using the DAGAN EX-1 amplifier, national instruments 6210 USB digitizer and WinEDR software. The following parameters were used: low-pass filtered at 3 kHz, high-pass filtered 0.3 Hz, digital gain 10 and sampling interval 10 μs. Single recordings were performed for ten minutes.

### Synthesis of MV1312 (3) and MV1369 (6)

Synthesis of MV1312 (4-Chloro-n-{3-[2-(4-methoxy-phenyl)-1h-imidazol-4-YL]-phenyl}-benzamide) and MV1369 (2-(3-methoxy-phenyl)-5-methyl-4-propyl-1h-imidazole) is depicted in **Scheme 1**. Details on the synthesis procedure can be found in the Supplementary data.

**Scheme 1.**
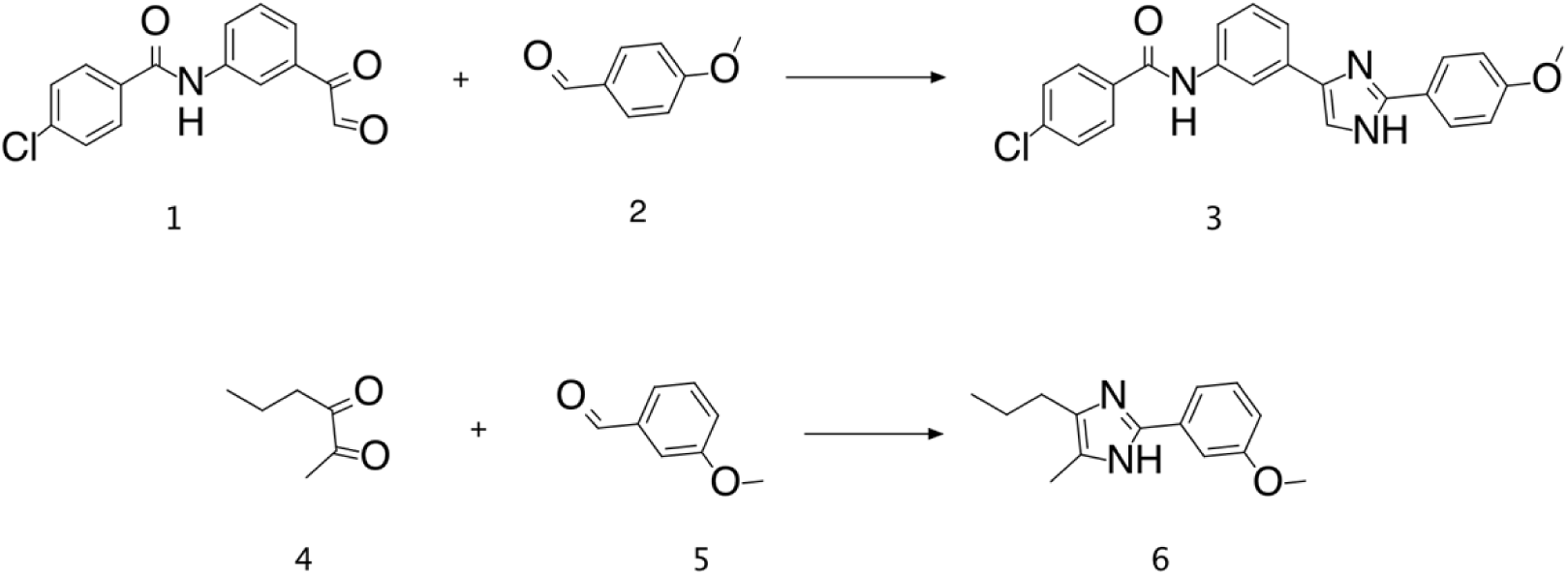
Reagents and conditions: 1 equiv of **2** and **5** with 4 equiv CH_3_COONH_4_ in 3.8 mL CH_3_OH; 1 equiv of **1** and **4** (respectively) in 3.5 mL CH_3_OH. Overnight rt.

### Determination of compound selectivity

Activity of MV1312 and MV1369 at hNav1.1–1.8 was assessed using a fluorescent imaging plate reader (FLIPRTetra, molecular devices) membrane potential assay as previously described^55^. In brief, cell lines (HEK293 Nav1.1–1.8) were plated on 384-well black-walled imaging plates (Corning) at a density of 10 000–15 0000 cells per well 48 hours before loading with 20 μ L of red membrane potential dye (proprietary formulation) (Molecular Devices, Sunnyvale, CA). Cells were incubated with the membrane potential dye for 30 min at 37 °C before the addition of compounds by the FLIPR^Tetra^ system. After the addition of 100µM MV1312, fluorescence was measured (excitation 515–545 nm; emission: 565–625 nm) for a period of 5 min to determine the effects of the compounds alone. Following this 5 min exposure, 5–20 μM VRT was added and fluorescence was measured for a further 5 min. Data was recorded and converted to response over baseline using Screenworks 3.2.0.14.

### Drug treatment

E3/drug solutions (2X) were prepared in 0.8% DMSO at their maximum tolerated concentration (MTC) as calculated elsewhere ^[17] [18] [33]^. In summary, the following MTCs were used: Carbamazepine (CBZ) 50µM, Phenytoin (PTH) 100µM, Valproic acid (VPA) 100µM, Stiripentol (STP) 12,5µM, Fenfluramine (FA) 50µM, Clemizole (CLM) 100µM. 500µL of AED solution was pipetted in each single well containing 500µL E3 generating a final DMSO concentration of 0.4%. For short incubation experiments embryos were incubated for 1 hour, for long incubation experiments, the incubation time was 18 hours. Compounds were ordered via Sigma-Aldrich.

### Determination of maximum tolerated concentration (MTC)

Compounds AA43279, MV1312 and MV1369 were tested for toxicity based on previously established methods ^[17]^ with slight modifications. In short; compounds were incubated in the bathing medium of 4 dpf larvae. After 24 hours, the following toxicity parameters were checked: touch response, loss of posture, body deformation and death. When none of these parameters were observed in any of the larvae tested, the concentration was regarded safe (data not shown).

### Statistical analysis

Data was analyzed and plotted using Graphpad Prism 7.04. Locomotor data did not pass the D’agostino & Pearson normality test, therefore the non-parametric Mann-Whitney U-test was used for data analysis. Mass-spectrometry was normally distributed and was further analyzed using the student t-test. Compound selectivity data was normally distributed and therefore analyzed with a multi-comparison ANOVA. A *P*-value of <0.05 was considered significant.

## Acknowledgement

The authors would like to thank Prof. Dr. J. Bakkers and Dr. F. Tessadori (Hubrecht Institute Utrecht, the Netherlands) for help and advice during the project. Compound AA43279 was a kind gift of Dr. T. Benned-Jensen (H.Lundbeck A/S Copenhagen). The authors would like to thank Prof. P. de Witte (K.U. Leuven, Belgium) and Dr. A. Zdebik (U.C. Londen, England) for their advice on the electrophysiology setup.

## Author contributions

Conceived project: BPCK KPJB. Designed and performed zebrafish experiments: WW SS MR RS MB NVD. Generated hSCN1A plasmid: LV. Designed and Performed compound synthesis and selectivity assays: MR MI IV. Analyzed the data and wrote the paper WW. MR. BPCK

